# The global expansion from southern Africa by modern humans? Its signal may be shown more through the cranial shape of males than females

**DOI:** 10.1101/2023.09.25.559024

**Authors:** Zarus Cenac

## Abstract

As the geographical distance between populations increases, populations are known to become more distant (dissimilar) in biological measures such as cranial shape. Relationships between geographical distance and biological distance are thought to be explained by modern humans having expanded from Africa. Expansion from Africa is also thought to explain why biological diversities (e.g., cranial shape diversity) decrease as geographical distance from Africa goes up. Research has suggested that modern humans expanded through different routes. If there were different routes, perhaps the relationship between geographical distance and cranial shape distance declines as the population (that distances are measured from) gets further away from Africa. This study used cranial shape measurements derived from the Howells data, and found that such a decline was suggested for males, unlike for females. It has been reasoned that variables which indicate the global expansion should signal an area of origin which is only in Africa. The relationship between geographical and cranial shape distances did not signal an exclusively African area. However, for males, the area was primarily in Africa. Overall, the relationship seemed to represent the expansion better amongst males than females. Congruently, cranial shape diversity declined more strongly for males than females as geographical distance from Africa accrued. To estimate which area of Africa the global expansion originated in, previous research pooled an estimate from genetic diversities, and suggested a southern origin. This estimate was supported when adding cranially-derived variables to the pooling, including the relationship between geographical distance and male cranial shape distance.

## Introduction

The biology of modern humans seems to provide clues as to how modern humans have swept across the world (von Cramon-Taubadel & Lycett, 2008). Globally, as the geographical distance between two populations rises, the genetic distance between the two populations greatens too (Ramachandran et al., 2005). Expansion from a common origin could lead to this pattern (Ramachandran et al., 2005). Examination of where an origin for a global expansion might be has pointed to Africa (e.g., Manica et al., 2007; Tishkoff et al., 2009). This global expansion could very well have begun in Africa 60-to-100 thousand years ago (Henn et al., 2012). Consideration has been granted as to whether bottlenecks were a feature of the expansion – with bottlenecks in expansion, genetic drift would increase (and diversity would decrease) the further a population is from the origin of the expansion (Ramachandran et al., 2005). Accordingly, interpopulation geographical and genetic distances are indeed related (Ramachandran et al., 2005), and the further away from Africa a population is located, the lower they are in several types of biological diversity (e.g., Balloux et al., 2009; Betti et al., 2009; Ramachandran et al., 2005) like cranial shape diversity (von Cramon-Taubadel & Lycett, 2008). As for which specific part/area of Africa possesses the origin of the expansion, there are a number of possibilities (e.g., Henn et al., 2012; Ray et al., 2005; Tishkoff et al., 2009) such as southwestern Africa (Tishkoff et al., 2009) or eastern (Ray et al., 2005).

### Distance and routes

Expansion from Africa of modern humans can explain not only genetic distance increasing with geographical distance (Ramachandran et al., 2005), but morphological distance increasing as well (Ponce de León et al., 2018). To be precise, with increasing geographical distance, genetic distance increases linearly (Lawson Handley et al., 2007; Ramachandran et al., 2005). Cranial distance also increases with geographical distance (Betti et al., 2010; Relethford, 2004a, 2009; von Cramon-Taubadel, 2019), although not at a constant rate, with cranial distance reaching towards a constant level (Relethford, 2004a) or apparently flattening out from a particular geographical distance (Betti et al., 2010).^1^ More specifically (see *Method*), interpopulation cranial *shape* distance rises with geographical distance (Hubbe et al., 2009).

It could be that modern humans expanded via different routes (Henn et al., 2012; Reyes-Centeno et al., 2014). For instance, research by Reyes-Centeno et al. (2014) involved geographical, genetic, and temporal bone (shape) distances, and they found support for modern humans having exited Africa and spread onward through a couple of routes – an earlier one which was relatively more southern, and a later one that was relatively more northern. Therefore, the relationship between geographical distance and biological distance could be expected to get weaker as the population (from which the geographical and biological distances are measured) gets further away from the origin of the expansion (Figure 1).

**Figure 1.**
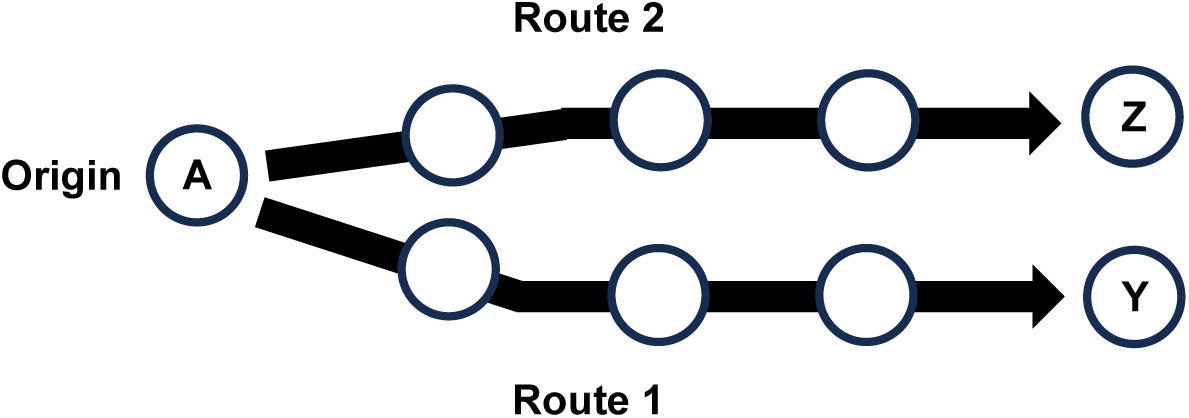
Routes in Expansion. *Note*. Figure 1 is based, to an extent, on an expansion scenario in Reyes-Centeno et al. (2014), and Figure 1 of Henn et al. (2012). Each circle is an example population, with Population A being the origin. Expansion is through bottlenecks, starting at the origin (e.g., Ramachandran et al., 2005) and going along two routes. Route 1 leads to Population Y, and Route 2 leads to Population Z. So, Populations Y and Z may be relatively close to each other geographically, but they arose from different routes (see Reyes-Centeno et al., 2014). The correlation between geographical and biological distances would be greatest when distances are from the population at the origin (Population A).

### Geographical area of origin and the strongest decline

It is possible to have an estimation of the geographical area in which the origin of the global expansion is located (e.g., Betti et al., 2009, 2013; Manica et al., 2007; Tishkoff et al., 2009). This can be achieved, for example, through seeing how much geographical distance from a place predicts diversity (e.g., Manica et al., 2007). For instance, the heterozygosity of autosomal microsatellites generates an area which is only in one continent, which is Africa (Manica et al., 2007). The area should only have geographical locations from which the diversity *decreases* most strongly – if the area is located 100% within Africa, then the measure (e.g., cranial shape diversity) may very well be indicative of the global expansion (e.g., Cenac, 2022b).

### Present study

This study gave some thought as to whether relationships between cranial shape and geographical distances are of value for locating the origin of the global expansion. Like with other variables (e.g., Cenac, 2022b, 2023), to infer whether the relationship between cranial shape and geographical distances indicates the global expansion from Africa, the present research looked not only for an indication of this relationship going through a decline as geographical distance rises from Africa, but for the relationship to decline the most when geographical distance is from a location in Africa rather than in another continent (i.e., for the suggested area of origin to not go beyond Africa). In some preprints, research has generalised across various measures to estimate the region of Africa where the global expansion started, and the region suggested was the south (Cenac, 2022b, 2023); if the relationship between cranial shape and geographical distances (Hubbe et al., 2009) is indicative of the global expansion, then this opens up the avenue of including this relationship alongside the various measures when generally estimating the origin.

## Method

Cranial shape measurements, calculated in previous research (Cenac, 2022a, 2022b), were used. These measurements (56 dimensions) were of Holocene populations of modern humans featured in the Howells dataset – 1,348 males (28 populations) and 1,156 females (26 populations), with males and females having 26 populations in common (Howells, 1973, 1989, 1995, 1996). The raw cranial data is at http://web.utk.edu/~auerbach/HOWL.htm. Cranial shape distances (as *D*-values) have been calculated before by using RMET 5.0 (Pinhasi & von Cramon-Taubadel, 2009); in the present study, RMET 5.0 (Relethford & Blangero, 1990) was used to calculate *D*^2^-values from cranial shape measurements, and the square root was taken of the *D*^2^ values to arrive at cranial shape distances (*D*-values). Geographical distances were from preceding research (Cenac, 2022a, 2022b) or found between coordinates (the coordinates being from Betti et al., 2013, and von Cramon-Taubadel and Lycett, 2008) using Williams (2011). Like beforehand (e.g., Cenac, 2022b), waypoints (Cenac, 2022a; von Cramon-Taubadel & Lycett, 2008) were used as appropriate when determining geographical distances. Other than RMET, calculations in the present research were undertaken in R Version 4.0.5 (R Core Team, 2021) and Microsoft Excel.

### Cranial shape distance

Rather than using interpopulation distances in cranial form (e.g., Betti et al., 2010; Roseman, 2004), the present study used cranial shape distances. The form of the cranium has size and shape facets (Betti et al., 2010). The relationship between cranial form and geographical distances could derive more from the shape of the cranium than its size (Cenac, 2022a). Indeed, with geographical distance, distance in cranial form increases (e.g., Relethford, 2004a) as does cranial shape distance (Hubbe et al., 2009). To the knowledge of the author, no study has seen if cranial size distance is associated with geographical distance. However, reference has been made to geographical distance being used as a stand-in for genetic distance (Relethford, 2013); whilst genetic distance increases with cranial shape distance, it does not increase with cranial size distance (Harvati & Weaver, 2006). Therefore, if the association between cranial distance and geographical distance does indicate the origin, a better indication of the origin may be found through cranial shape distances than through cranial form distances. Moreover, the global expansion may be expressed better through cranial shape diversity than through cranial form diversity (Cenac, 2022b). And so, cranial shape distance was utilised over cranial form distance. Following Betti et al. (2010), heritability was set to 1.00 when calculating distances, a value which was also used in von Cramon-Taubadel and Pinhasi (2011).

### Spearman correlation coefficients

For the relationships between cranial shape distance and geographical distance, specifically Spearman correlation coefficients were calculated. Why is this? The association between genetic distance (using autosomal microsatellites) and geographical distance follows a linear trend (Lawson Handley et al., 2007; Ramachandran et al., 2005) as mentioned in the *Introduction*. However, the relationship between cranial and geographical distances seems to be embodied better with a non-linear trend than a linear one (Betti et al., 2010).^2^ The author of the present study was unaware of any preceding study having determined if there is a non-linear relationship between cranial shape distance and geographical distance. Nonetheless, if the association between cranial form and geographical distances (e.g., Relethford, 2004a) does particularly arise from the relationship between cranial shape and geographical distances (Cenac, 2022a), then non-linearity likely would be present. It is not known if the relationship between cranial and geographical distances would have the same non-linear pattern across all populations (from which distances are measured), or indeed if the non-linear pattern would be driven by only some populations. So, conservatively, Spearman correlation coefficients were used in this study for the relationship between cranial shape and geographical distances.

### Origin of the global expansion

Various locations within Africa (99 locations) (Betti et al., 2013) and worldwide (32 locations) (von Cramon-Taubadel & Lycett, 2008) have been utilised as possible origins (Cenac, 2022b),^3^ and were used as such in the present study.

Earlier research has utilised the Bayesian information criterion (BIC) to signal the geographical area which the global expansion started in (e.g., Manica et al., 2007). BICs were used for that purpose in the current study. In that methodology regarding BICs and expansion, the area of origin is made of locations – the location with which the relationship between some variable and geographical distance (to populations) gives the *lowest* BIC, and other locations with which relationships give rise to BICs which the lowest BIC is within four BICs of (Manica et al., 2007). The direction of relationships is of importance – the *lowest* BIC (and other BICs in the area of origin) should be in a particular direction (e.g., Cenac, 2022b). As with the association between autosomal diversity and geographical distance from the origin (e.g., Ramachandran et al., 2005), the origin in the present study should be signified by a decline – the association between the relationship between cranial shape and geographical distances ought to decline (rather than increase) as geographical distance from the origin grows (see above). So, as with genetic diversity (e.g., Cenac, 2022b), the location generating the lowest BIC, and the remaining locations forming the area of origin, were defined from locations which produce declines. BIC calculation used a formula that is featured in Masson (2011). In that formula, BIC is calculated from *R*^2^ (Masson, 2011). Regarding the formula, squared *r*_s_ has been used in the place of *R*^2^(Cenac, 2022b), and was used as such in the present study for female crania when estimating the area of origin (concerning nonparametric analysis).

Correlations between coefficients and geographical distance from each of several locations in Africa were found and ranked (like Cenac, 2023, did for correlations between genetic diversity and geographical distance from Africa) in order to see what contribution results in the present study would make to the broad estimate Cenac (2023) found for the part of Africa which the global expansion originated in.

### Figures

The design of graphs displaying correlation coefficients, BICs, and distances (including continental symbols) carried on from previous research (Cenac, 2022a, 2022b). This pertains to Figures 2, 3, and 6. A new addition is the inclusion of red lines to indicate the geographical locations with BICs which are at values between the lowest BIC and (within) four BICs over the lowest. Figure 4 is, to some extent, based on Figure 4 of Cenac (2023) (who employed origins that are in a figure in Betti et al., 2013). Figure 5 modifies Figure 11A from Cenac (2022b), being produced from the relationship between cranial shape and geographical distances rather than (as in Cenac, 2022b) cranial shape diversity.

**Figure 2.**
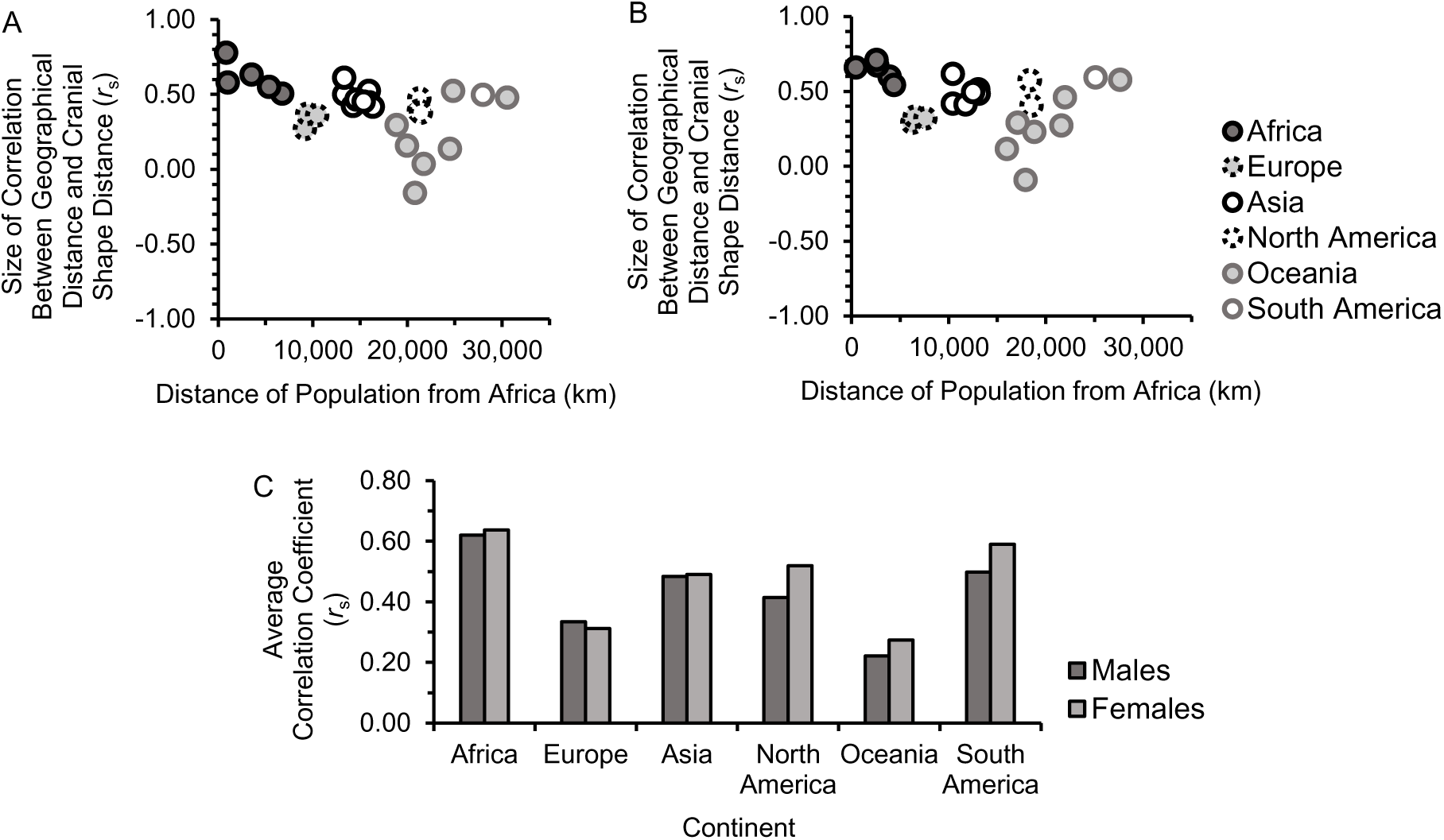
Cranial Shape and Geographical Distances: Decline from Africa? *Note*. Each datapoint in Figure 2A and 2B is for the correlation between two variables – i) cranial shape distance from a population, and ii) geographical distance from that same population. Figure 2A is for male crania, whilst 2B is for female crania. On the *y*-axes of 2A and 2B, the datapoint is placed according to how far the population (which the distances are from) is located geographically from the place which generated the *lowest* BIC. For Figure 2C, correlation coefficients were averaged within the continental categories shown in 2A and 2B. Coefficients can be Fisher *z*-transformed, mean averaged, and means converted to coefficients (Corey et al., 1998) – that procedure was used for Figure 2C. Continents are ordered in Figure 2C (and in the legend for 2A and 2B) based on averaging (mean) population distances from Africa; continents are placed in ascending order of averaged geographical distance from Africa.

**Figure 3.**
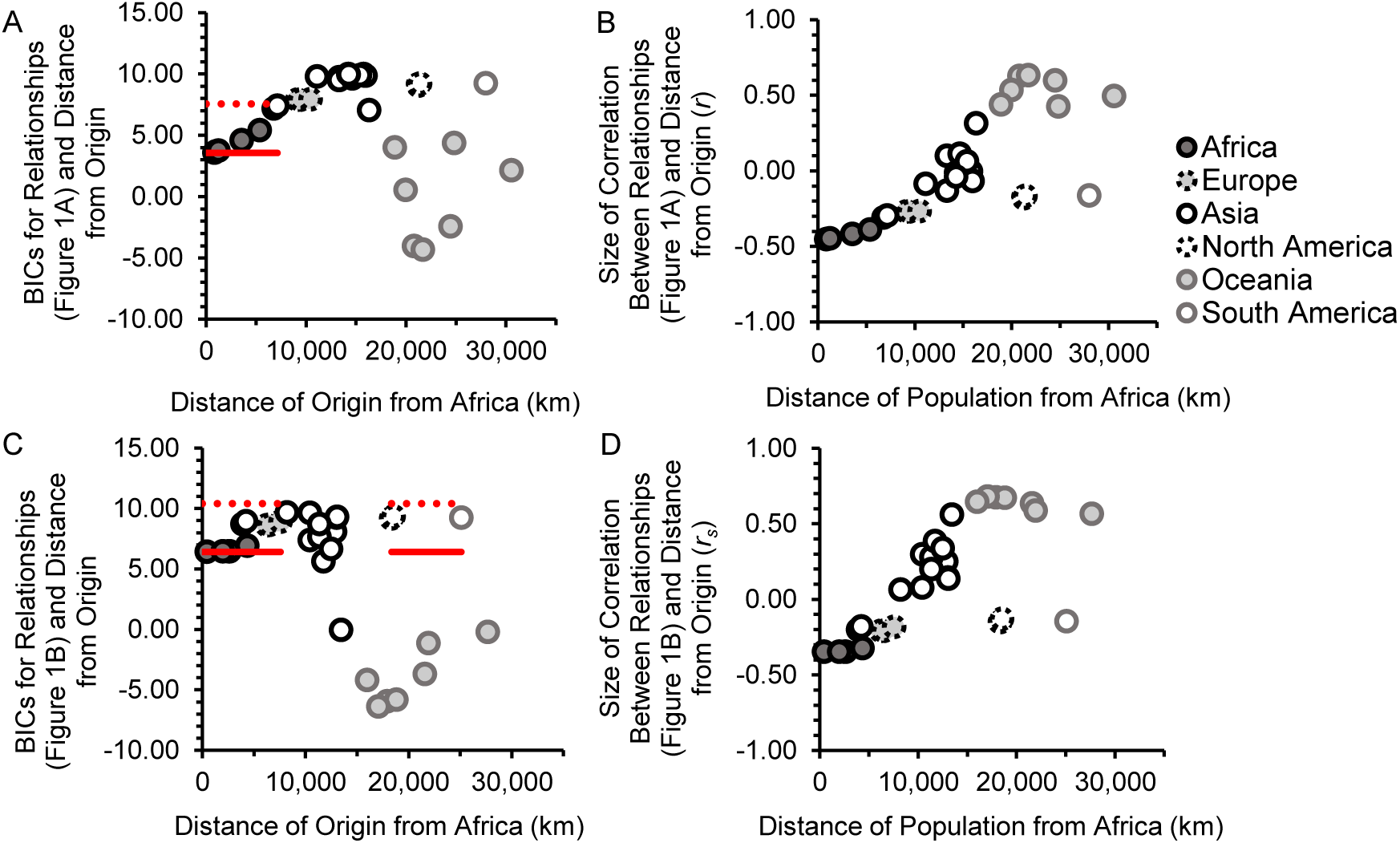
Declines and Origins. *Note*. Continents are ordered in the legend of Figure 3 the same as in Figure 2. Distances on *y*-axes are from the geographical location which had the lowest BIC (of locations which produced negative correlation coefficients). Horizontal red lines represent the lowest BIC (solid red line) and four BICs above it (the dotted red line). Although circles (BICs, *r*s, and *r*ss) are only shown for 32 locations, BICs, *r*s, and *r*ss were also calculated for a further 99 locations which were each in Africa (the locations in Betti et al., 2013), and the *lowest* BIC was found to be amongst the 99 for males and females.

**Figure 4.**
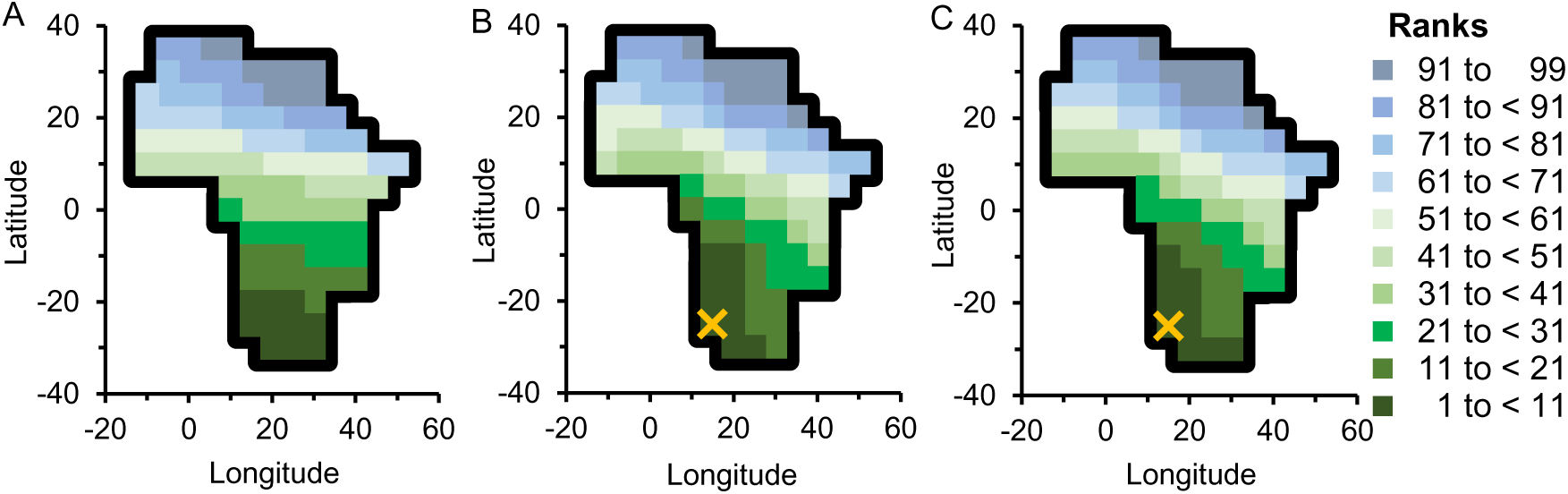
Origin in Southern Africa is Supported. *Note*. For Figure 4A, correlation coefficients were found measuring the association between two variables. Variable 1 is the relationship between cranial shape and geographical distances for males (the coefficients in Figure 2A). Variable 2 is the geographical distance between an origin in Africa and each of the populations (which distances are measured from in Variable 1). So, between Variables 1 and 2, a correlation coefficient was found using each African origin. The strengths of these associations were ranked as in Cenac (2023), with a rank of number 1 for the most negative, and these ranks are presented in Figure 4A. It should be reiterated that the area of origin was not 100% in Africa (see Figure 3A). For Figure 4B, ranks calculated in the presented study for the relationship between cranial shape and geographical distances were pooled with the ranks in Cenac (2023) for types of genetic diversity (autosomal, mitochondrial, X-chromosomal, and Y-chromosomal). Lastly, in Figure 4C, ranks calculated regarding cranial shape diversity and cranial sexual size dimorphism were pooled with the ranks used in 4B. In Figure 4B and 4C, the X is the location with the highest rank (the rank of number 1).

**Figure 5.**
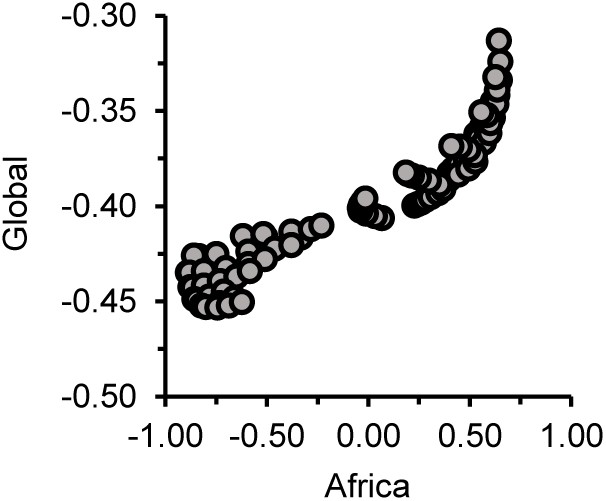
Ranked Correlation Coefficients (Figure 4A) are Likely due to Distances from Populations in Africa. *Note*. In Figure 5, each datapoint is a correlation coefficient (*r*) representing when a location in Africa was used as an origin. So, each datapoint represents the correlation between distance from that origin and the relationship between cranial shape and geographical distances. On the *x*-axis of Figure 5, correlation coefficients were found when using distances from African populations to other populations whether these other populations are in Africa or elsewhere. On the *y*-axis, distances were from each of the 28 populations. As noted in Cenac (2022b) regarding that research, there would be nonindependence.

**Figure 6.**
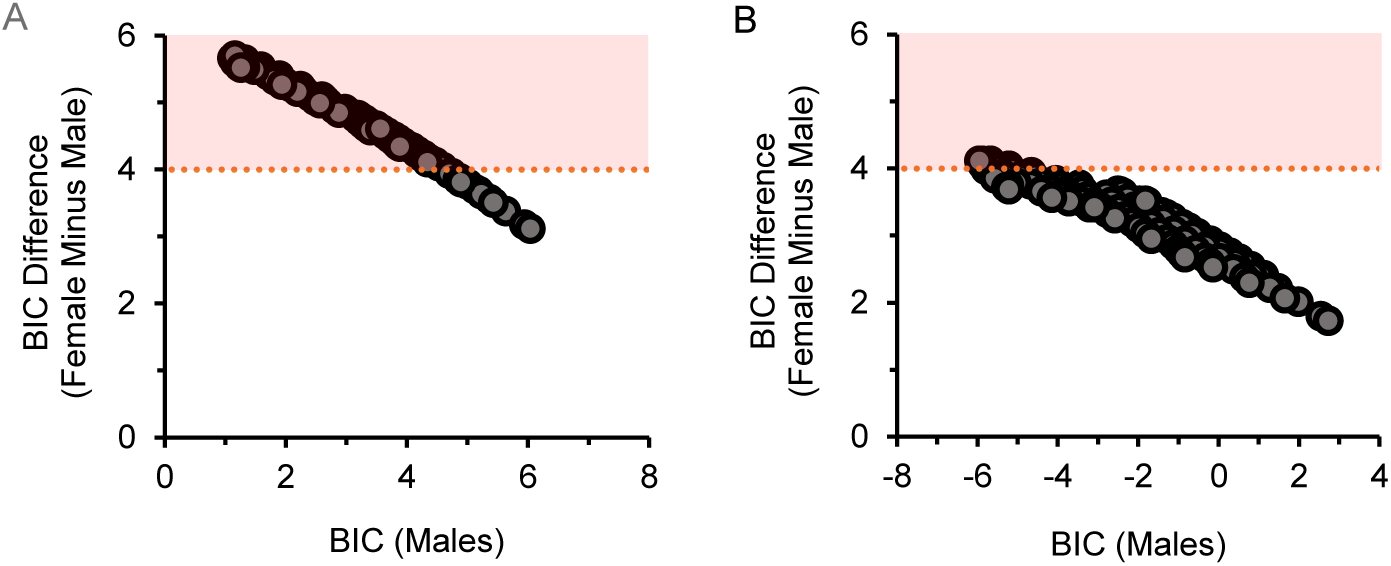
Choice of Origin may Affect Whether a Difference in Relationships with Geographical Distance from Africa is Found (Males Compared to Females) *Note*. Figure 6A pertains to some of the BICs for the analysis of whether the relationship between cranial shape and geographical distances is associated with geographical distance from Africa. Figure 6B is with respect to BICs for the analysis regarding the relationship between the shape diversity of crania and geographical distance from Africa. BICs in the red area are for geographical distances from Africa which lead to a notable difference (using the aforementioned four-BIC technique, e.g., Manica et al., 2007).

### Correlation and nonindependence

When examining if geographical distance from Africa and diversity show a relationship, a way of picking the African place (which the geographical distances are from) uses the location that results in the lowest BIC (Atkinson, 2011). In the current study, that method was used regarding testing if a correlation happens between geographical distance from Africa and the relationship between cranial shape and geographical distances. The lowest BIC was also used when comparing the strengths of associations (see *Results and discussion*).

As with the association which diversity has with geographical distance from Africa (Balloux et al., 2009; Cenac, 2022b, 2023; Prugnolle et al., 2005; von Cramon-Taubadel & Lycett, 2008), it would be useful to use *p*-values to infer whether the strength of relationships (between cranial shape and geographical distances) is related to geographical distance from Africa. Yet, nonindependence was probably great enough to be an issue because the same populations contributed towards multiple correlation coefficients (see Forstmeier et al., 2017). It is not clear how the potential nonindependence in the present study could be resolved. It is known that nonindependence can lead to *p*-values being underestimated (Forstmeier et al., 2017); *p*-values were nonetheless calculated in the present study, but they are (therefore) by no means definitive, and yet if an incorrect *p*-value suggests a non-significant result then it is therefore likely that the correct *p*-value would signal non-significance.

Holm-Bonferroni adjustments (Holm, 1979) were utilised on *p*-values. These adjustments were done using the Gaetano (2013) spreadsheet. To check if residuals (parametric analysis) were spatially autocorrelated, a spreadsheet from Chen (2016) was used which produces a spatial Durbin-Watson (*DW*). It is acceptable to use bounds for the Durbin-Watson statistic on the spatial *DW* (Chen, 2016) – 5% bounds (Savin & White, 1977) were indeed used for that purpose in the present study. Of several ways of defining atypical datapoints, residuals were standardised by transforming them into *z*-scores, with transformed values greater than |3.29| being defined as atypical (e.g., Field, 2013).

## Results and discussion

### Decline?

For males, cranial shape and geographical distances might very well become less related as the population (that distances are from) gets geographically further away from Africa, *r*(26) = -.45, *p* = .031, spatial *DW* = 1.61 (Figure 2A). However, as mentioned above, nonindependence is likely to have been at play. Although, visually, a decline seems to be indicated (Figure 2A), South America does seem to diverge from the possible trend (Figure 2A and 2C). However, only one population in the Howells data is located in South America (Howells, 1989) (see Figure 2A), so the correlation coefficient found for South America should particularly be regarded with some uncertainty.

With respect to female crania, the distribution of correlation coefficients (at distances from the location with the lowest BIC) was indicated to have heteroscedasticity. Therefore, a nonparametric analysis was used for female crania; a decline was not indicated, *r*_s_(24) = -.35, *p* = .081 (Figure 2B)^4^ – as there probably was nonindependence, the *p*-value likely would be underestimated (see Forstmeier et al., 2017), which suggests that there was indeed no significant decline.

### Strongest decline?

For males, BICs suggested an area of origin which covered all of the African origins used in the present study. This area was not in Africa alone (Figure 3A). Of the origins used outside of Africa, the area of origin featured only one location. That location was for the Tel Aviv coordinates from von Cramon-Taubadel and Lycett (2008), which gave a BIC that was 3.84 BICs above the lowest, and was therefore only just within the area of origin (Figure 3A). Therefore, the area of origin encompassed Africa, and was only slightly in Asia. And so, for males, the area of origin is largely (although not fully) consistent with a global expansion from Africa, with the strongest declines generally being from Africa.

Regarding the global expansion, the relationship between cranial shape and geographical distances for males appears to be reminiscent of hip bone shape diversity in Betti et al. (2013). Betti et al. (2013) observed that diversity in hip bone shape exhibited a decrease with the greatening of geographical distance from Africa, and hip bone shape diversity showed an area of origin, the majority of which was in Africa. Still, in Betti et al. (2013), hip bone shape diversity still seems to have been regarded like it is suggestive of global expansion from Africa. And so, like hip bone shape diversity (Betti et al., 2013), an argument could be made for the global expansion being indicated through the relationship between male cranial shape distance and geographical distance.

As for female crania, the area of origin did not at all particularly indicate an African origin (Figure 3C and 3D), which might seem unsurprising given the lack of correlation. Indeed, there seemed to be at least two areas, neither of which only featured Africa. Actually, BICs were of no use in determining an area of origin with female crania, with the area being defined wholly on the basis of whichever direction correlation coefficients were in (Figure 3C and 3D).

Therefore, results with males are broadly consistent with the global expansion from Africa being indicated in the relationship between cranial shape and geographical distances. Yet those results are not entirely consistent given that some proportion of the area of origin is not within Africa. As for females, the relationship would not seem to be sufficiently indicative of the global expansion.

In some preprints, there have been attempts to interpolate between estimates of origins from several biological variables in order to have a broad impression of which part of Africa the expansion originated, with the most plausible region appearing to be southern Africa (Cenac, 2022b, 2023). This has been done, for example in Cenac (2023), through ranking locations in Africa in terms of how well geographical distances from those locations are related to a variable (given a certain direction), and then pooling together those ranks from a number of variables in order to have an overall evaluation of where the origin is likely to be located. It seems that southern Africa (Choudhury et al., 2021) is given some further support by the ranks generated in the present study regarding the relationship between cranial and geographical distances concerning male crania (Figure 4A), including when these ranks are used to supplement the pooled estimate in Cenac (2023) (Figure 4B).^5^

### Difference in results for males and females

This study has suggested that the relationship between cranial shape and geographical distances could very well be a useful indicator of the global expansion when male crania are used, but not when female crania are utilised. Perhaps this should have been foreseen because, in Cenac (2022a), some indication may have been found for distances within cranial dimensions, as a product of the global expansion, being larger amongst males than females.

In the present study, whilst 26 populations were used for analysis with female crania, an additional two populations were used when analysing male crania. Could the inclusion of the additional two populations have been a factor in the expansion likely being indicated through male, but not female, crania? To answer this question, analysis with the male crania was redone utilising just the 26 populations in the Howells (e.g., 1989, 1996) data that have both males and females. Using the 26 populations, the decline in the relationship between cranial shape and geographical distances went from *r* = -.45 with the 28 populations to *r* = -.53 with the 26 populations (the lowest BIC was at the same location whether using 28 or 26 populations). Moreover, the area of origin became smaller when using 26 populations, with the area no longer being in both Africa and Asia, but now only being in Africa. And so, whilst males had different results to females, this difference would not seem to have been because of the additional two populations being included in the analysis with males.

When calculating the area of origin, for male crania, BICs were calculated from squared Pearson correlation coefficients. However, for female crania, BICs were calculated from squared Spearman coefficients. When squared Pearson coefficients were used to calculate BICs and define the area(s) of origin using female crania, the areas were the same as the ones indicated in Figure 3C. These areas were also designated on the basis of whether coefficients were negative.

Furthermore, the different results do not seem to be due to the relationship between cranial shape and geographical distances being greater for males than females, because the relationship actually does not appear to be greater for males (Table 1). Therefore, the difference in results between males and females may be due to something related to the global expansion.

**Table 1.** The Relationship Between Cranial Shape Distances and Geographical Distances. *Note*. It had already been found that geographical distance explains cranial form distance in non-linear analysis at *R*^2^ = 33% for males and also for females too (Betti et al., 2010); as for cranial shape, in the present study, the relationship between geographical distance and cranial shape distance was numerically higher for females than males.

| Subset of the Howells data | Correlation coefficient ($r_s$) |
| --- | --- |
| Males (28 populations) | .38 |
| Males (26 populations) | .36 |
| Females (26 populations) | .42 |

Additionally, when applying BICs in the context of the expansion (whether BICs are inside four BICs of whatever the lowest BIC is) (e.g., Manica et al., 2007), the lowest BIC for males (the 26 populations) was substantially lower than the lowest BIC for females. And so, there was a difference between males and females in terms of i) whether there was a significant decline (Figure 2), ii) whether Africa was generally (selectively) indicated as the origin (Figure 3), and also iii) the strength to which the relationship between cranial shape and geographical distances is associated with geographical distance from Africa. Therefore, it appears that the relationship between cranial shape and geographical distances is a better reflection of the global expansion amongst male crania than amongst female crania.

#### Cranial shape diversity

Expansion from Africa by modern humans gives an explanation for the relationship between biological and geographical distances (Ponce de León et al., 2018; Ramachandran et al., 2005). It also provides an explanation for the decreasing of biological diversity with the accumulation of geographical distance from Africa (e.g., Ramachandran et al., 2005; von Cramon-Taubadel & Lycett, 2008). For instance, the global expansion explains why a decrease in cranial shape diversity is found as geographical distance from Africa increases, and this decrease happens for male crania (von Cramon-Taubadel & Lycett, 2008) and female crania (Cenac, 2022b). Therefore, results in the current study might lead to the expectation of the decrease of cranial shape diversity being stronger for males than for females. Indeed, in previous research, the decrease (as distance from Botswana increased) was observed to be bigger *numerically* for males than females in terms of correlation coefficients (Cenac, 2022a). This led to the prospect of the expansion being more evident though cranial shape diversity amongst males than females (Cenac, 2022a). Yet, it was untested whether there was a notable difference in the correlation coefficients (Cenac, 2022a). In the present study, the decline of cranial shape diversity (with the greatening of geographical distance from Africa) was found to be notably greater for males than females (Table 2), which seems consistent with the global expansion indeed being exhibited more in the cranial shape diversity of males than females. Hence, the difference between males and females in the decline of cranial shape diversity (Table 2) appears to be agreeable with findings in the present research regarding the relationship between cranial shape and geographical distances.

**Table 2.** Cranial Shape Diversity: Stronger Expansion Signal for Males than for Females. *Note*. Cranial shape measurements had been calculated using the Howells data (26 populations of modern humans) beforehand (Cenac, 2022a, 2022b); using those measurements, the cranial shape diversity of the 26 populations was found for males and females in RMET 5.0.^6^ The geographical distance giving the lowest BIC (of distances giving a negative correlation coefficient) was distance from San. This happened for males and females also.^7^ BICs were used to choose the best models regarding expansion – being within a certain number of BICs (four) of the lowest of the BICs (e.g., Manica et al., 2007). Geographical distance from San provided a BIC for males which was more than four BICs lower than the BIC which geographical distance from San gave for females. Therefore, the global expansion seemed to have a stronger signal in cranial shape diversity for males than females.

| Subset of the Howells data | Correlation between diversity and geographical distance from San (*r*) |
| --- | --- |
| Males (26 populations) | -.67 |
| Females (26 populations) | -.60 |

### Origin: Southern Africa

Both cranial shape diversity and cranial sexual size dimorphism appear to be indicative of the global expansion (Cenac, 2022b). The pooling in Cenac (2023) only used genetic diversities, rather than cranial variables. This was due to uncertainty over how to include cranial variables (see Cenac, 2023, for details); in the present study, cranial variables were included in the pooling. The cranial shape diversity of males is indicated to reflect the global expansion more than the cranial shape diversity of females (Table 2). Therefore, male cranial shape diversity (rather than female cranial shape diversity) was included in the pooling. These diversities were for the 28 populations in the Howells data, and the diversities used were calculated in Cenac (2022b) (although, cranial shape diversity was calculated in von Cramon-Taubadel & Lycett, 2008). The pooling also included cranial sexual size dimorphism. The dimorphism used in this study was calculated in Cenac (2022a) for Howells data populations which have males and females – the present study used dimorphism for populations, and these populations were ones used in correlational analysis in Cenac (2022b) (24 populations), who excluded two populations to deal with positive spatial autocorrelation.

As geographical distance from Africa increases, there is a decline in cranial shape diversity (Cenac, 2022b; von Cramon-Taubadel & Lycett, 2008) and an increase in cranial sexual size dimorphism (Cenac, 2022b). And so, origins were ranked higher for shape diversity according to how negative correlation coefficients are (the most negative being ranked first) as Cenac (2023) did regarding genetic diversities, and ranked higher for size dimorphism according to how positive the coefficients are (the most positive being ranked first).

Following/extending Cenac (2023), for various origins in Africa (99 places), an average of ranks was taken at each origin for autosomal diversity, mitochondrial diversity, X-chromosomal diversity, Y-chromosomal diversity, the relationship between cranial shape and geographical distances (males), cranial shape diversity (males), and cranial sexual size dimorphism. Also following/extending that study (Cenac, 2023), these averaged ranks were then ranked (across origins). These ranks (across origins) are shown in Figure 4C. Southern Africa (Choudhury et al., 2021) is indicated by the ranks as being where the global expansion started (Figure 4C).

### Distance from San

For males, and females, San (Howells, 1989, 1996) was the population from which the relationship between cranial shape and geographical distances seemed to be at its (numerically) most positive point (Figure 2A and 2B). Out of populations in the Howells (1973, 1989, 1995, 1996) data, it may be expected that the relationship would reach its most positive when distances were from San if the relationship actually reflects the global expansion. This can be said based on population divergences (Li et al., 2008; Schlebusch et al., 2020) and diversity levels (Balloux et al., 2009; Cenac, 2022b; Colonna et al., 2011; Hunley et al., 2012; Manica et al., 2007; Prugnolle et al., 2005; Ramachandran et al., 2005; Schlebusch et al., 2020; von Cramon-Taubadel & Lycett, 2008).

Regarding divergences amongst modern humans, the (numerically) earliest divergence (mean time) appears between Khoe-San and other populations (Schlebusch et al., 2020; but see Skoglund et al., 2017). In a phylogenetic tree, divergences between populations do seem to broadly be congruent with expansion from Africa (Li et al., 2008). In that study, which used 51 HGDP-CEPH populations, the divergence most near to the root was between San and other Africans (Li et al., 2008). In a global expansion featuring bottlenecks, populations nearer to where the expansion originated would have undergone less drift and be more diverse (Ramachandran et al., 2005). Several types of genetic diversity are related to geographical distance, with these diversities falling with increasing geographical distance from Africa (Balloux et al., 2009; Prugnolle et al., 2005) in particular (Cenac, 2022b; Manica et al., 2007; Ramachandran et al., 2005). Autosomal heterozygosity has been found to be higher for Khoe-San than for other Africans (Schlebusch et al., 2020). Regarding the Howells cranial data (28 populations for males, and, for females, 26 populations), diversity in cranial shape declines the further from Africa (in particular) populations are, and the numerically highest cranial shape diversity is found to be with San (Cenac, 2022b; von Cramon-Taubadel & Lycett, 2008).

However, San, in the Howells data, were from a considerable geographical expanse, which could enhance the diversity in their crania (Algee-Hewitt, 2011). For males, the relationship between cranial shape and geographical distances appears to generally be congruent with the global expansion from Africa, and this congruency may therefore be given further credence by the relationship being at its numerically most positive when distances are from San.

### Distance from Africa

If locations inside of the African continent are treated as origins, African populations may predominantly determine how relatively well distance from an origin relates to cranial shape diversity (Cenac, 2022b). This may also seem to happen if cranial shape diversity is replaced with the relationship between male cranial shape and geographical distances (Figure 5). Therefore, the ranks shown in Figure 4A are likely to be largely attributable to the cranial and geographical distances from populations in Africa (to other populations regardless of whether they are also in Africa or outside).

The African continent has five of the Howells data populations (Howells, 1989, 1996; von Cramon-Taubadel & Lycett, 2008). Do distances from these five populations really result in a pattern which is indicative of the expansion? Studies have been in line with not only there being expansion from Africa (e.g., Ramachandran et al., 2005), but movement back into the continent (e.g., Hodgson et al., 2014) and admixing (e.g., Gallego Llorente et al., 2015), with Eurasian ancestry having a presence in the continent (e.g., Haber et al., 2016; Pickrell et al., 2014; Tishkoff et al., 2009). For instance, Dogon are part of the Howells data (e.g., Howells, 1989, 1996), and Dogon have been found to have 44.5% European or Middle Eastern ancestry, yet that figure was calculated for a sample of just nine individuals (Tishkoff et al., 2009). Nonetheless, in a dendrogram constructed from the Howells data, Dogon do cluster with other sub-Saharan Africans (Relethford, 2009). The Egyptian population in the Howells data is suggested to have a phenotypical affinity with populations outside of Africa (Algee-Hewitt, 2011). Indeed, the Egyptian population (Howells data) does seem to be reflective of Greek ancestry to some extent (although, six cranial dimensions were used in that study – six were common across the data used) (Sanders et al., 2014). Research is consistent with admixture affecting genetic distance (Ramachandran et al., 2005). For males, the relationship between cranial shape and geographical distances seems to be greater for Africans than others (Figure 2A and 2C). And so, Eurasian ancestry within Africa could have contributed to Figure 4A.^8^

### Future research

Ideally, research will continue considering whether the global expansion is exhibited more through the morphology of males than females. The relationship between hip bone shape diversity and geographical distance from Africa has seemed suggestive of being stronger for males (27 populations) than females (20 populations), although not when the same populations (19) are used for males and females (Betti et al., 2013). That research, which was Betti et al. (2013), used the Akaike information criterion (AIC) to estimate the area of origin with respect to hip bone shape diversity. The area of origin was made of the location that brought about the lowest AIC, and locations that gave AICs within four AICs of that lowest one (Betti et al., 2013). When exploring whether the decline in hip bone shape diversity differs between males and females, Betti et al. used geographical distances from the centroid of the area of origin. The centroid was not at the location which gave the lowest AIC for either the males from the 27 populations or the females from the 20 populations (Betti et al., 2013). Betti et al. (2013) do not seem to have stated whether the centroid would be the same as the location which would give the lowest AIC for the 19 populations (which featured both males and females). For the analyses in the present study with cranial shape and geographical distances, and cranial shape diversity, it does seem that finding a difference between males and females (in relationships with geographical distance from Africa) was facilitated by using geographical distance from the African location which had the lowest BIC (Figure 6). Therefore, it could be worth seeing whether the decline of hip bone shape diversity (as geographical distance from Africa ascends) is stronger for males than females specifically when using the lowest AIC (if this has not been done yet).

Particularly for the male crania, a remaining concern is the possible nonindependence. Aside from cranial shape distance (Hubbe et al., 2009), other biological distances are related to geographical distance (e.g., Betti et al., 2014; Ramachandran et al., 2005). It would be useful to determine whether such relationships indicate the origin of the global expansion by applying the methodology which was utilised in the present study. Finding them to be indicative would support the soundness of using the relationship between biological and geographical distances to estimate the origin of the global expansion. This would go some way to lessening concerns about the likely nonindependence.

## Conclusion

The relationship between biological and geographical distances could be useful when seeking the origin of the global expansion. Usefulness was broadly indicated in the present study when biological distances were of the cranial shape of males. When pooled with other variables, the relationship between cranial shape and geographical distances (males) added further weight to the global expansion having its origin in southern Africa. Different routes in expansion from Africa (Henn et al., 2012; Reyes-Centeno et al., 2014) could be responsible for the global expansion likely being indicated through the relationship between the cranial shape of males and geographical distances. Interpopulation distance in cranial shape seems to reflect the global expansion more so for male crania than female crania. This does appear to be backed up by results regarding within-population diversity in cranial shape. It is not clear why the expansion signal seems to appear more for males than females in cranial shape.

## Footnotes

1 Genetic similarity (estimated through crania) is related to geographical distance, and this could be explained by expansion from Africa (Relethford, 2004b); genetic distance estimated via crania (e.g., Relethford, 2013) can be regarded as cranial distance (e.g., Pinhasi & von Cramon-Taubadel, 2009; von Cramon-Taubadel & Pinhasi, 2011).

2 The association between cranial form and geographical distances was explored using populations globally in Relethford (2004a, 2009) and Betti et al. (2010). Relethford (2004a, 2009) used the Howells dataset, with 26 populations being employed by Relethford (2004a), whereas Relethford (2009) utilised 18 populations. Betti et al. (2010) used 105 and 39 populations for males and females respectively. A greater extent of flattening appears in Betti et al. (2010) than in Relethford (2004a, 2009). Unlike Betti et al. (2010), Relethford (2004a, 2009) did not do some sort of comparison between the fit of linear and non-linear models.

3 Betti et al. (2013) used those 99 African locations as origins (and others), and von Cramon-Taubadel and Lycett (2008) used four of the worldwide locations as origins (plus others also).

4 For female crania, four locations had the lowest BIC. To choose which one to use as the location giving the lowest BIC (in the correlation test, and on *y*-axes in Figures 2B, 3C, and 3D), each of the four locations was given a number from 1 to 100, and the one with the lowest number was selected as the one treated as having the lowest BIC.

5 The pooling method used in Cenac (2023) was extended to also feature ranks calculated in the present study (from correlation coefficients measuring how much the relationship between cranial shape and geographical distances is associated with geographical distance from Africa) alongside ranks calculated in Cenac (2023). Cenac (2023) used ranks for several diversities to arrive at a pooled estimate of the origin – autosomal (microsatellite, and single nucleotide polymorphism haplotype), mitochondrial, X-chromosomal, and Y-chromosomal. The autosomal diversities were sourced from Pemberton et al. (2013) (microsatellite) and Balloux et al. (2009) (single nucleotide polymorphism haplotype), the mitochondrial and X-chromosomal diversities were sourced from Balloux et al. (2009) too, and the Y-chromosomal diversity was sourced from de Filippo et al. (2011) (Cenac, 2023). Balloux et al. (2009) used autosomal haplotype diversity from Li et al. (2008). See those previous studies for details on where the data used to calculate the diversities were themselves sourced from (Balloux et al., 2009; de Filippo et al., 2011; Li et al., 2008; Pemberton et al., 2013).

6 Using the Howells data, cranial shape diversity for the 26 populations had been calculated previously – separately for males and females (Cenac, 2022a).

7 For females in the 26 populations, BICs had been calculated for the relationship between cranial shape diversity and geographical distance (Cenac, 2022b), and they were recalculated in the current study.

8 For autosomal diversity, it has previously been reasoned that non-African ancestry could have some influence when estimating the origin of global expansion (Cenac, 2022b). Nonetheless, the autosomal diversity (microsatellite heterozygosity) of sub-Saharan Africans suggests that the global expansion originated in the southern part of Africa, with this suggestion being found regardless of whether analyses adjusted diversity (of sub-Saharan Africans) for the non-African ancestry present in populations (Cenac, 2022b).

